# Factors of sex and age dictate the regulation of GABAergic activity by corticotropin-releasing factor receptor 1 in the medial sub-nucleus of the central amygdala

**DOI:** 10.1101/2020.07.22.215947

**Authors:** Siara Kate Rouzer, Marvin R. Diaz

**Affiliations:** Department of Psychology, Center for Development and Behavioral Neuroscience, Binghamton University, Binghamton, NY 13902, United States; Developmental Exposure Alcohol Research Center, Binghamton University, Binghamton, NY 13902, United States

**Keywords:** corticotrophin-releasing factor, age, sex, central amygdala, GABA, development

## Abstract

Adolescents are phenotypically characterized with hyper-sensitivity to stress and inappropriate response to stress-inducing events. Despite behavioral distinctions from adults, investigations of developmental shifts in the function of stress peptide corticotrophin-releasing factor (CRF) are generally limited. Rodent models have determined that CRF receptor 1 (CRFR1) activation within the central amygdala is associated with a stress response and induces increased GABAergic synaptic neurotransmission within adult males. To investigate age-specific function of this system, we performed whole-cell patch clamp electrophysiology in brain slices from naive adolescent (postnatal days (P) 40-49) and adult (>P70) male and female Sprague Dawley rats to assess GABAergic activity in the medial central amygdala (CeM). Our results indicate a dynamic influence of age and sex on neuronal excitability within this region, as well as basal spontaneous and miniature (m) inhibitory post-synaptic currents (IPSCs) in the CeM. In addition to replicating prior findings of CRFR1-regulated increases in mIPSC frequency in adult males, we found that the selective CRFR1 agonist, Stressin-1, *attenuated* mIPSC frequency in adolescent males, at a concentration that did not affect adult males. Importantly, this age-specific distinction was absent in females, as Stressin-1 attenuated mIPSC frequency in both adolescent and adult females. Finally, only adult males exhibited an increase in mIPSC frequency in response to the CRF1R antagonist, NBI 35965, suggestive of tonic CRFR1 activation in the CeM of adult males. Together, these data emphasize the robust influence of age and sex on neurophysiological function of a brain region involved in the production of the stress response.

## Introduction

Stress is an experience that produces both protective and harmful outcomes (McEwen, 1998). When stress surpasses an intrinsic threshold, the sympathetic nervous system is activated, setting off a chain-reaction of neurochemical events that trigger negative behavioral and psychological consequences (see review: (Klein and Corwin, 2002))). Across species, adolescents demonstrate heightened vulnerability to stress compared to adults. In humans, adolescents exhibit elevated activity in areas associated with emotional reactivity, such as the amygdala, following exposure to threat (fearful faces) (Hare et al., 2008). Adolescent rats (Postnatal Days [P]28-50) demonstrate 1) elevated corticosterone (CORT) response to stressors and 2) a prolonged return to baseline CORT levels compared to adult counterparts (Brown and Spencer, 2013). This enhanced activity within the hypothalamus–pituitary–adrenal (HPA) axis has been similarly observed in adolescent primate and avian models, and appears to wane by adulthood (Andersen and Teicher, 2008). This age-specific activity is hypothesized to involve extra-hypothalamic regions which continue to develop in adolescence, although the exact mechanisms are still under investigation.

Sex has also been shown to dictate behavioral and neural responses to stress, primarily in adults. For instance, sex can determine whether a stressor enhances or diminishes cognitive performance in fear-learning assessments (Brown and Spencer, 2013). However, the direction and degree of stress-induced response in females is still widely contended. We and others have shown that compared to males, females demonstrate sex-specific resilience to the negative effects of stress in both hamsters (Faruzzi et al., 2005) and rats (Bowman et al., 2003; Varlinskaya et al., 2020), an effect which was blocked in rats with administration of an estrogen antagonist (Wood and Shors, 1998). However, several independent studies have also reported that females exhibit greater stress-induced HPA activity compared to males (Kudielka and Kirschbaum, 2005; Walker and McCormick, 2009; Young et al., 2007). Notably, sex-differences in HPA and autonomic responses appear to be most prominent when females were assessed within a pubertal-menopausal window (Kajantie and Phillips, 2006). This suggests that age as a biological factor contributes to stress reactivity.

The central amygdala (CeA) is a region well-characterized for its role in regulating stress and anxiety-like responses. This striatal-like structure is composed primarily of GABAergic local interneurons and projection neurons (Babaev et al., 2018; Ehrlich et al., 2009). Inhibitory tone within the medial subnucleus of the CeA (CeM) has been associated with reduced response to stress-inducing stimuli (Gilpin et al., 2015), possibly through direct efferents to brain regions responsible for the expression of anxious behaviors (Babaev et al., 2018). This region is also rich in concentrations of the stress peptide corticotrophin-releasing factor (CRF). CRF is a 41 amino-acid peptide found in many mammalian species, and is identical across human and rat species (Dautzenberg and Hauger, 2002). Throughout the brain, high concentrations of CRF and activation of one of its primary receptors, CRFR1, have been recognized for their anxiogenic influence in adult populations (Kehne, 2007; Magalhaes et al., 2010).

Within the CeM of adult male rodents, CRF potentiates GABA release through presynaptic CRFR1, and selective antagonism of CeM-CRFR1 blunts anxiety-like behavior (Skorzewska et al., 2017). However, response to CRFR1 activation in adolescents is largely unexplored. One investigation of presumed-adolescent male Sprague Dawley rats (125-150g) has demonstrated that CRF superfusion into the CeA results in neuronal hyperpolarization (Rainnie et al., 1992), however, this study did not directly examine synaptic transmission. A more recent study in the lateral habenula showed a CRFR1-induced reduction in GABA transmission in post-weanling rats (Authement et al., 2018), *opposite* of what has been established in the adults within the CeA (Bajo et al., 2008; Nie et al., 2004; Roberto et al., 2010). These findings suggest a potential developmental shift in CRFR1-regulation of GABA transmission, however age and sex as biological factors have not been systematically investigated.

Thus, the objective of the present study was to determine if the biological factors of age and sex influence CRF system function within the CeM. Using whole cell electrophysiology, we determined that these factors significantly influence basal GABA transmission, as well as the instrinsic excitability of CeM neurons. Furthermore, CRFR1-regulated GABAergic transmission within this region was age and sex-specific in both the sensitivity of presynaptic CRFR1 and the direction of receptor-regulated GABA release. Finally, we determined that adult males, and not adolescent males or adolesent/adult females, demonstrate tonic CRFR1 activity.

## Materials and Methods

### Animals

To avoid introducing shipping stress during development, all experimental subjects were bred in-house as previously described. Adult male and female Sprague Dawley breeders were obtained from Envigo/Harlan (Indianapolis, IN) and permitted to acclimate at least one week prior to breeding. Upon detection of sperm in vaginal smears, pregnant dams were isolated and housed with a plastic hut and crinkle paper as nesting material. After parturition, litters were culled to 5:5 males/females on P2 and housing conditions remained the same with a plastic hut and crinkle paper. Pups were weaned from their mother on P21 and housed with same-sex littermates until experimentation. All animals were group-housed (2-3 animals per cage) in a temperature-controlled (22°C) vivarium and maintained on a 12:12 h light:dark cycle (lights on at 7:00 h). Subjects were provided *ad libitum* access to food (5L0D PicoLab Laboratory Rodent diet) and water throughout the duration of experimentation. All animal procedures were approved by the Binghamton University Institutional Animal Care and Use Committee.

### Whole-cell Electrophysiology

#### Drugs and Chemicals

All chemicals and kynurenic acid were purchased from Sigma-Aldrich (St. Louis, MO, USA). APV, tetrodotoxin (TTX), Stressin-1, CGP 55845 and NBI 35965 were purchased from Tocris/R&D systems (Bristol, UK).

#### Slice Preparation

Adolescent (P40-48) and adult (P70-95) male and female rats were sedated with a 250 mg/kg dose of ketamine and quickly decapitated, as previously performed by our lab (Przybysz et al., 2017). Brains were rapidly removed and immersed in ice-cold oxygenated (95% O_2_-5% CO_2_) sucrose artificial cerebrospinal fluid (ACSF) containing (in mM): sucrose (220), KCl (2), NaH_2_PO_4_ (1.25), NaHCO_3_ (26), glucose (10), MgSO_4_ (12), CaCl_2_ (0.2), and ketamine (0.43). Coronal slices (300 μM) containing the CeM were collected with a Vibratome (Leica Microsystems. Bannocknurn, IL, USA). Slices were incubated in 34°C normal ACSF (in mM): NaCl (125), KCl (2), NaH_2_PO_4_ (1.3), NaHCO_3_ (26), glucose (10), CaCl_2_ (0.1), MgSO_4_ (0.1), ascorbic acid (0.04), and continuously bubbled at 95% O_2_-5% CO_2_ for at least 40 minutes before recording. All experiments were performed within 6 hours of slice preparation.

#### Whole-cell patch-clamp recordings

Following incubation, slices were transferred to a recording chamber, where oxygenated ACSF was warmed to 32°C and continuously superfused over the submerged slice at 3.3 ml/min. Recording electrodes of 3-5 MΩ tip resistance were pulled from borosilicate glass capillary tubing (Sutter Instruments) using a Flaming-Brown puller (Sutter Instruments). Recordings were collected from the CeM with patch pipettes filled with experiment-specific internal solutions (see below). Data were acquired with a MultiClamp 700B (Molecular Devices, Sunnyvale, CA) at 10 kHz, filtered at 1 kHz, and stored for later analysis using pClamp 10 software (Molecular Devices).

#### Voltage-clamp recordings

For voltage-clamp experiments, a KCl-based internal solution was used for detecting GABA_A_ receptor-mediated currents, containing (in mM): KCl (135), HEPES (10), MgCl_2_ (2), EGTA (0.5), Mg-ATP (5), Na-GTP (1), and QX314-Cl (1); 300 mOsm; 7.3 pH with KOH. For recordings of GABA_A_ receptor-mediated spontaneous inhibitory postsynaptic currents (sIPSCs), AMPA and NMDA glutamate receptors were pharmacologically blocked using 1 mM kynurenic acid and 50 μM APV, respectively, and GABA_B_ receptors were blocked with CGP 36645 (1 μM). For data collection of miniature IPSCs (mIPSCs), recordings were made in the presence of the Na+ channel blocker, TTX (1 μM). Baseline recordings of sIPSCs and mIPSCs were allowed to equilibrate for at least 5 min before recording began. A 3-min baseline period was recorded prior to drug application.

For electrophysiological assessment of CRFR1 function, the CRFR1 selective-agonist, Stressin-1 (10 nM, 100 nM and 1 μM) or the selective antagonist, NBI 35965 (1 μM), was applied for 15 min. As interneurons within the CeM exhibited generally stable access resistance, this length of time ensured that changes in transmission could be visualized for stability. To determine the timing of a stable drug effect, a time course of drug exposure was constructed during the pilot phase of experimentation and stable activity was observed 5 min following drug application. Therefore, the first 5 min of drug application was removed from final analyses. Additionally, for all experiments, only recordings with an access resistance change of <20% are included in final analyses.

#### Current-clamp recordings

For assessments of intrinsic excitability and resting membrane potential (RMP), recordings were collected from the CeM with patch pipettes filled with a K-gluconate-based internal solution containing (in mM): K-Gluconate (120), KCl (15), EGTA (0.1), HEPES (10), MgCl_2_ (4.0), MgATP (4.0), Na_3_GTP (0.3) and Phosphocreatine (7.0); 280-290 mOsm; 7.25 pH with KOH. Cells were opened in a voltage-clamp configuration (holding potential = −70 mV) and switched to current-clamp settings for at least 7 minutes to allow neurons to dialyze prior to applying a series of depolarizing current steps (15 pA, 500 ms duration). Subsequent firing activity was recorded for future analysis. Rheobase, action potential threshold and time to first action potential peak were determined with the first action potential fired with the lowest stimulation current as assessed by pClamp 10 software (Molecular Devices).

### Experimental Design and Statistical Analysis

Based on preliminary data, a power analysis (*G*Power 3.1.9.2*) for a two-way (sex x age) analysis of variance (ANOVA) and an alpha of 0.05 suggested a sample size of 8 units per group. The unit of analyses for all experiments was cell, with no more than 2 cells per animal used for any given experiment. Furthermore, no more than 2 subjects per litter were used for any given experiment to reduce litter effects in our sample population. To avoid experimenter bias, all data analyses were conducted by an individual blind to the conditions of the subject. All statistical analyses were performed using GraphPad 8 Software (Prism). Measures of membrane properties, RMP, intrinsic excitability, and basal s/mIPSCs were analyzed with 2 (age: adolescents, adults) x 2 (sex: male, female) between-subjects ANOVA. Upon discovering a significant sex x age interaction in basal sIPSC frequency in the CeM, all CRFR1-targetted analyses were subsequently analyzed independently in each sex. Concentration-response activity following CRFR1 activation were reported as % change from baseline mIPSC activity and analyzed independently in each sex using 2 (age) x 3 (concentration: 10 nM, 100 nM, 1 μM) between-subjects ANOVAs. CRFR1 activity was measured in each age/sex group by performing a one-sample *t-test*, statistically comparing the average change in mIPSC baseline activity to a null mIPSC change (0). Significance was defined as *p* ≤ 0.05 in all assessments. In the event of significant main effects or interactions, *post hoc* Sidak’s multiple comparison tests were performed to determine specific group differences. All data are presented as mean ± standard error of the mean (SEM) unless otherwise specified.

## Results

### Membrane properties of CeM interneurons do not differ across age and sex

GABAergic interneurons in the CeM were identified by their approximate membrane resistance to membrane capacitance values, as previously reported (Herman et al., 2016). Assessment of membrane properties across age (adolescent and adult) and sex (male and female) revealed no significant group differences in membrane capacitance [age: *F*(1,60) = 0.758, *p* = 0.388; sex: *F*(1,60) = 2.085, *p* = 0.154; age x sex interaction: *F*(1,60) = 0.028, *p* = 0.868, *n* = 12-20 cells per group] or membrane resistance [age: *F*(1,60) = 0.163, *p* = 0.689; sex: *F*(1,60) = 2.455, *p* = 0.122; age x sex interaction: *F*(1,60) = 0.003, *p* = 0.955, *n* = 12-20 cells per group] (*Table 1*).

**Table 1.** Cell properties of CeM interneurons across age and sex, reported as Mean (SEM). Experimental groups do not differ in native membrane properties or resting membrane potential. However, adults demonstrate generally lower action potential thresholds than adolescents, independent of sex. Both sex and age influence time to action potential peak following current injection, with adults and females demonstrating the quickest times. * indicates significant main effect of age (p < 0.05), # indicates significant main effect of sex (p < 0.05).

| | Membrane Resistance (MΩ) | Membrane Capacitance (pF) | Resting Membrane Potential (mV) | AP Threshold (mV) * # | Rheobase (pA) | Time to Peak (ms) * |
| --- | --- | --- | --- | --- | --- | --- |
| Adolescent Males (n=14) | 424.21 (70.00) | 41.66 (4.56) | -51.59 (2.80) | -45.55 (2.41) | 52.50 (7.16) | 123.75 (15.36) |
| Adolescent Females (n=18) | 529.05 (82.78) | 46.69 (4.37) | -48.99 (1.86) | -43.40 (1.31) | 55.00 (8.47) | 101.58 (22.28) |
| Adult Males (n=16) | 392.32 (41.70) | 37.57 (2.43) | -51.63 (1.77) | -50.07 (1.70) | 50.00 (6.12) | 73.18 (18.78) |
| Adult Females (n=18) | 504.98 (56.00) | 43.92 (3.38) | -49.76 (2.10) | -45.54 (1.12) | 56.36 (9.37) | 67.96 (12.94) |

### Both sex and age influence the excitability of CeM neurons

In a current-neutral configuration, cells were assessed for resting membrane potential. Neither age nor sex influenced resting membrane potential [age: *F*(1,62) = 0.037, *p* = 0.849; sex: *F*(1,62) = 1.104, *p* = 0.297; age x sex interaction: *F*(1,62) = 0.029, *p* = 0.865, *n* = 14-18 cells per group] (*Table 1*). Cells were then held at −70 mV to assess and normalize differences in excitability across groups. Notably, the proportion of cells that demonstrated firing activity following current injection were considerably lower in adolescent males compared to adolescent females (Males: 6/14, Females: 14/18) but comparable between sexes in adulthood (Males: 10/16, Females: 11/18).

From cells that were responsive to current injection, we found no differences in rheobase across ages and sexes [age: *F*(1,36) = 2.109, *p* = 0.155; sex: *F*(1,36) = 0.041, *p* = 0.840; age x sex interaction: *F*(1,36) = 0.077, *p* = 0.784] (*Table 1*). However, age did significantly influence the membrane potential at first AP [*F*(1,36) = 4.356, *p* = 0.044], with adults demonstrating lower thresholds than adolescents, an effect driven by adult males (*Table 1*). Furthermore, females demonstrated significantly higher thresholds than males [*F*(1,36) = 4.403, *p* = 0.043]. There was no significant interaction between these two variables [*F*(1,36) = 0.555, *p* = 0.461]. Quantification of firing activity revealed a significant age x sex interaction [*F*(1,703) = 19.48, *p* < 0.001], whereupon adolescent males exhibited significantly greater activity than adult males [*t*(1,23) = 3.300, *p* = 0.0031] (*Fig. 1A*). This effect of age was absent in females [*t*(1,32) = 0.962, *p* = 0.343] (*Fig. 1C*). We also found a significant main effect of age when examining time to first AP peak, with adults demonstrating quicker onset of the first action potential than adolescents [*F*(1,37) = 4.141, *p* = 0.049] (*Table 1* and depicted in exemplar traces in *Fig. 1B&D*). This activity was not influenced by sex, either as a main effect [*F*(1,37) = 0.438, *p* = 0.512] or as an interacting variable [*F*(1,37) = 0.168, *p* = 0.684].

**Figure 1.**
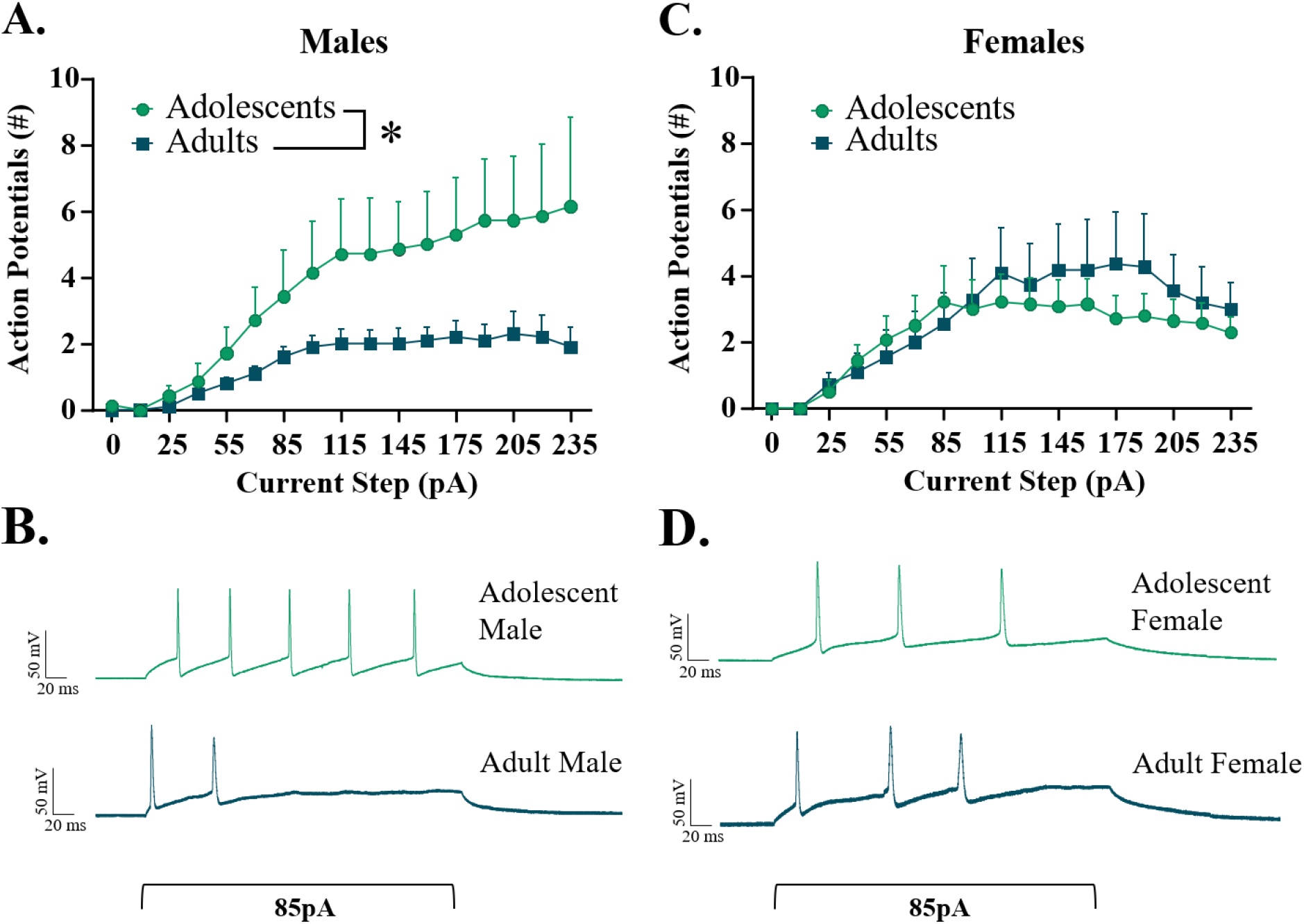
Excitability of interneurons within the CeM, Males (A&B) and Females (C&D). Increasing current steps (pA) produced significantly more firing activity in adolescent males than adult males, with no difference in females between adolescent and adult age groups. * indicates significant difference between groups (p < 0.05).

### Basal synaptic transmission in the CeM is sex- and age-specific

To determine if factors of age and sex influenced basal inhibitory synaptic activity in the CeM, both sIPSCs and mIPSCs were recorded. As represented in *Fig. 2A*, analyses of basal sIPSC frequency revealed a significant interaction of age x sex [*F*(1,66) = 6.296, *p* = 0.014] (*Fig. 2B*), with no main effects of age [*F*(1,66) = 1.830, *p* = 0.181] or sex [*F*(1,66) = 1.720, *p* = 0.194]. Post-hoc analyses revealed significantly higher sIPSC frequency in adolescent females compared to both age-matched males (*n* = 14-20, *p* = 0.048), and adult females (*n* = 16-20, *p* = 0.035). Notably, males did not differ in sIPSC frequency between age groups (*n* = 14-20, *p* = 0.853). Assessment of sIPSC amplitude revealed a significant main effect of sex [*F*(1,67) = 10.590, *p* = 0.002], with females consistently exhibiting greater post-synaptic receptor amplitude than males (*Fig. 2C*), while age did not influence sIPSC amplitude (age: [*F*(1,67) = 0.016, *p* = 0.900]; age x sex interaction: [*F*(1,67) = 0.4737, *p* = 0.494]).

**Figure 2.**
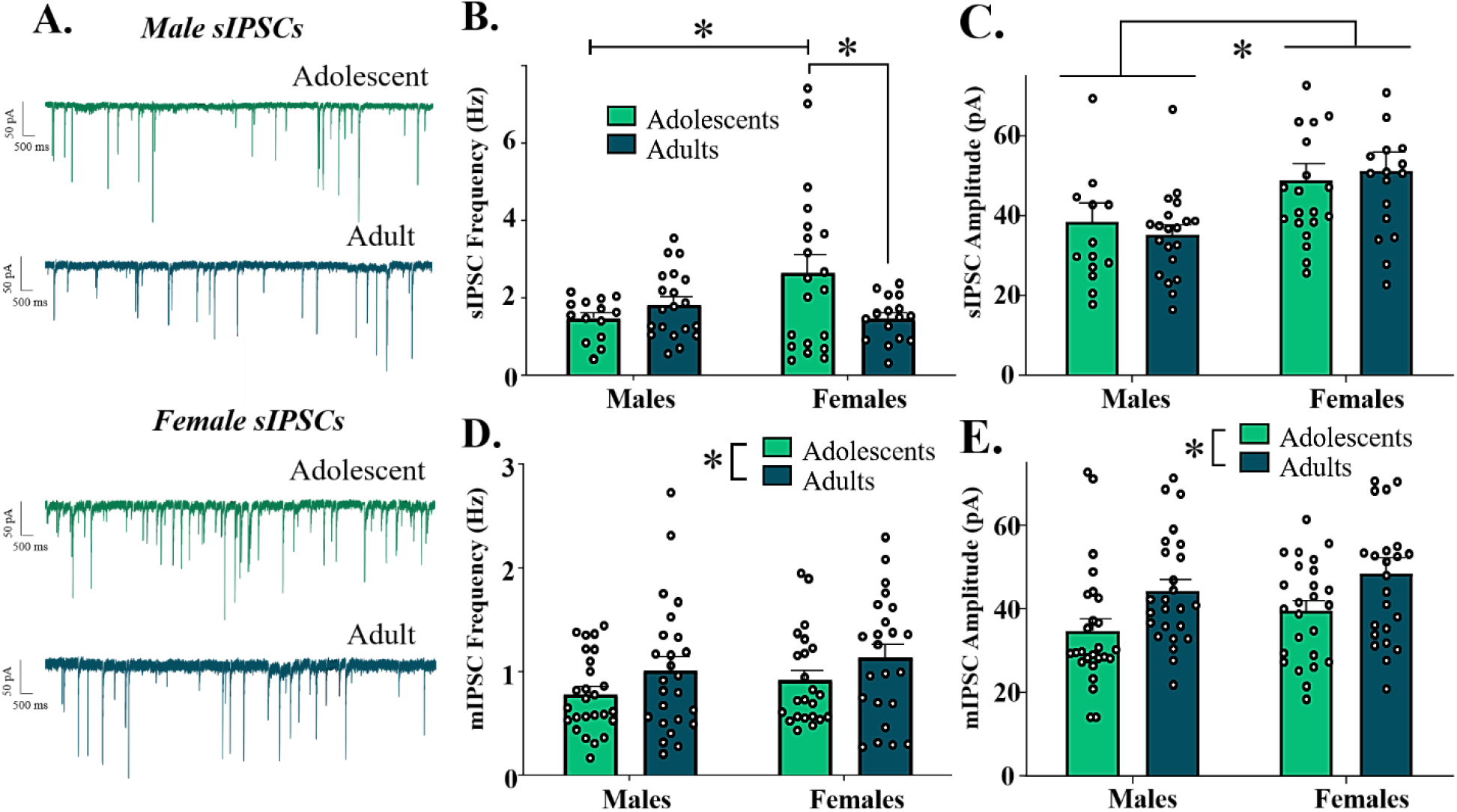
Basal IPSCs in the CeM across age and sex. (A) Representative sIPSC activity of neurons in the CeM. B) sIPSC frequency across experimental groups. Adolescent females exhibited significantly greater sIPSC frequency than both adolescent males and adult females. C) sIPSC amplitude across experimental groups. Females exhibited significantly higher sIPSC amplitude than males, independent of age. D) mIPSC frequency across experimental groups. Adults demonstrated greater mIPSC frequency than adolescents, independent of sex. E) mIPSC amplitude across experimental groups. Adults demonstrated higher mIPSC amplitude than adolescents, independent of sex. * indicates significant difference between groups (p < 0.05)

In contrast, action potential-independent mIPSC frequency significantly differed only between age groups [*F*(1,93) = 4.559, *p* = 0.035], with adults exhibiting higher mIPSC frequency than adolescents (*Fig. 2D*). This effect was not significantly different between sexes (main effect of sex: [*F*(1,93) = 1.461, *p* = 0.223]; age x sex interaction: [*F*(1,93) = 0.002, *p* = 0.964]). A similar pattern emerged in assessment of mIPSC amplitude, with adults demonstrating greater amplitude than adolescents [*F*(1,93) = 10.49, *p* = 0.002] (*Fig. 2E*), independent of sex (main effect of sex: [*F*(1,93) = 2.462, *p* = 0.120; age x sex interaction: [*F*(1,93) = 0.010, *p* = 0.924]).

### CRFR1 activation bi-directionally regulates GABAergic transmission in adolescent and adult males

Given the age x sex interactions found in basal sIPSC activity, CRFR1-regulated activity was subsequently analyzed independently in each sex. Previous research in adult, male Sprague Dawley rats has shown that CRFR1 activation within the CeA increases mIPSC frequency without changing mIPSC amplitude (Herman et al., 2013a; Kang-Park et al., 2015; Roberto et al., 2010). To determine if the function of CRFR1 on GABA transmission is age-dependent, we assessed the effect of the CRFR1-selective agonist, Stressin-1 (10 nM, 100 nM and 1 μM) on mIPSCs within the CeM of adolescent and adult males.

Analysis of Stressin-1-induced changes in mIPSC frequency revealed significant main effects of both age [*F*(1,43) = 30.700, *p* < 0.001] and concentration of drug [*F*(2,43) = 4.364, *p* = 0.019] in males, and a significant age x dose interaction [*F*(2,43) = 8.320, *p* < 0.001], with a leftward shift in concentration in adolescent males (*Fig. 3A*). Post-hoc analyses revealed no significant difference between adolescents and adults at the 10 nM concentration [*t*(1,43) = 0.066, *p* > 0.999], whereupon 10 nM Stressin-1 did not produce a significant change in mIPSC frequency in either adolescents [*t*(1,7) = 0.164, *p* = 0.875, *n* = 8 cells] or adults [*t*(1,7) = 0.056, *p* = 0.957, *n* = 8 cells]. At 100 nM, drug effects were significantly different between age groups [*t*(1,43) = 4.294, *p* < 0.001]; this concentration produced no significant change in mIPSC frequency in adults [*t*(1,7) = 0.218, *p* = 0.884, *n* = 8 cells], however 100 nM Stressin-1a significantly reduced mIPSC frequency in adolescents [*t*(1,9) = 8.642, *p* < 0.001, *n* = 10 cells]. Finally, at 1 μM, a significant difference between age groups was also observed [*t*(1,43) = 5.356, *p* < 0.001]. At this concentration, Stressin-1 continued to significantly reduce mIPSC frequency in adolescents [*t*(1,7) = 7.089, *p* < 0.001, *n* = 8 cells], yet produced an opposite and significant increase in mIPSC frequency in adults [*t*(1,7) = 9.576, *p* < 0.001, *n* = 8 cells] (*Fig. 3B*).

**Figure 3.**
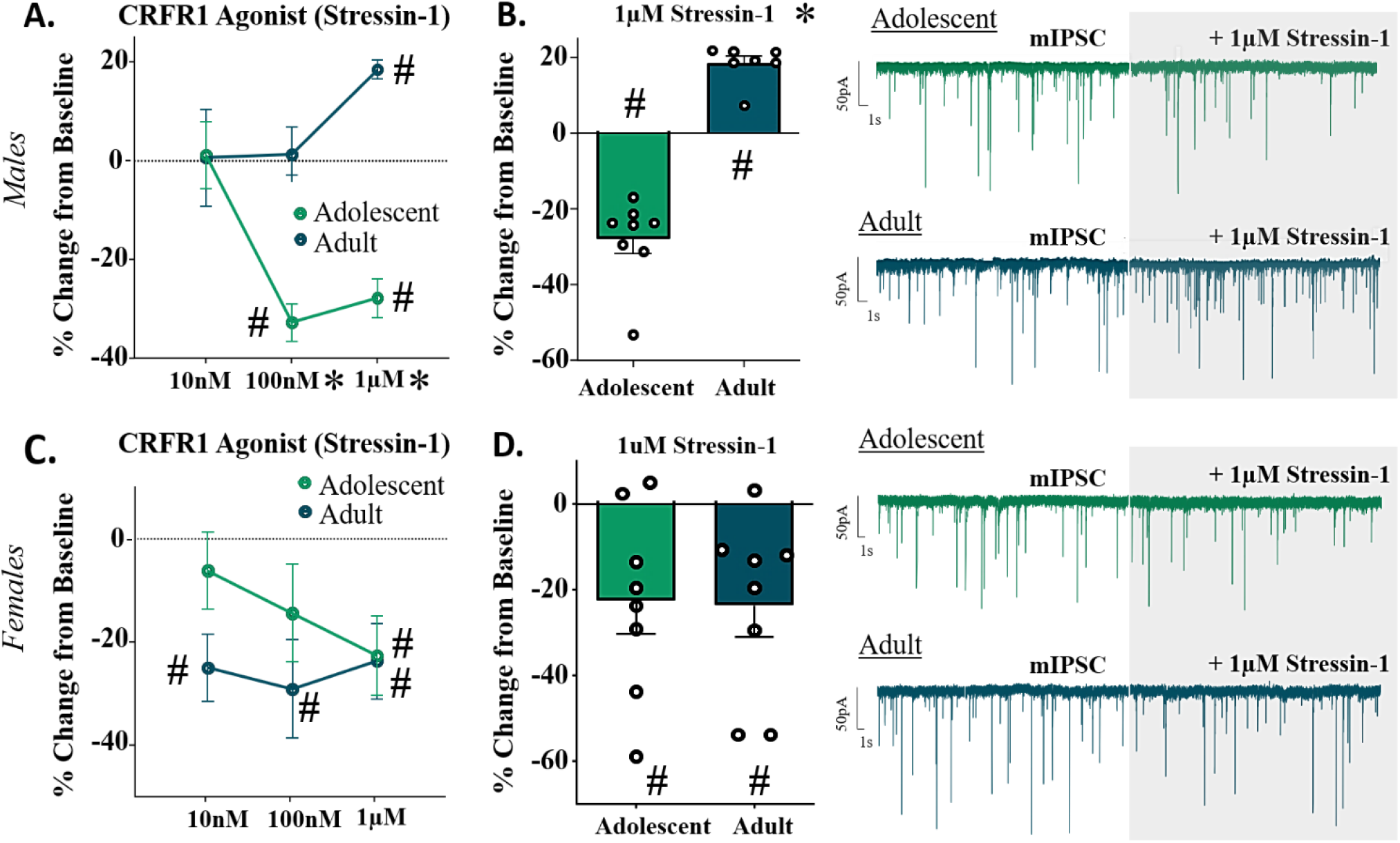
Change in mIPSC Frequency (%) following bath application of selective CRFR1-receptor agonist, Stressin-1. A) In males, CRFR1 activation produces significant dose-dependent changes in both adolescents and adults. In adolescents, 100nM and 1μM Stressin-1 significantly decrease mIPSC frequency, while 1μM Stressin-1 produces a significant increase in mIPSC frequency in adults. B) Bar graph and representative traces of % change in mIPSC frequency from adolescent and adult males following bath application of 1μM Stressin-1. C) In females, CRFR1 activation produces significant reductions in mIPSC frequency. In adolescents, 1μM Stressin-1 significantly decreases mIPSC frequency, while all three doses of Stressin-1 produce a significant decrease in mIPSC frequency in adults. D) Bar graph and representative traces of % change in mIPSC frequency from adolescent and adult females following bath application of 1μM Stressin-1. * indicates significant effect of age (p < 0.05). # signifies significant difference from 0.

Analysis of drug-induced changes in mIPSC amplitude in these same cells revealed no significant main effects of age [*F*(1,43) = 2.172, *p* = 0.148] or dose of drug [*F*(2,43) = 0.255, *p* = 0.776] in males, nor a significant age x drug interaction [*F*(2,43) = 0.884, *p* = 0.421] (*Table 2*).

**Table 2.**
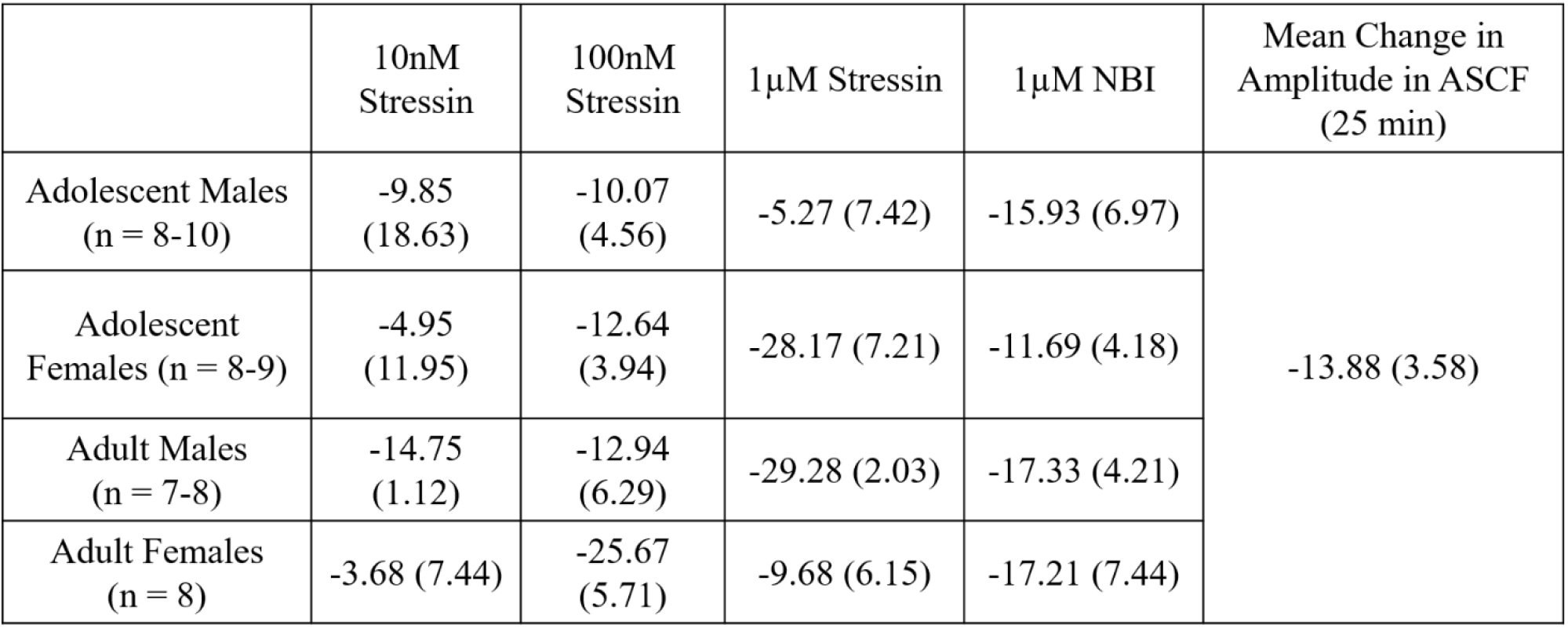
% Change in mIPSC amplitude per experimental group for each CRFR1-targeted experiment. There were no significant effects of sex or age on mIPSC amplitude at any dose/drug. Far-right column represents the average decline in postsynaptic receptor amplitude in a subset of drug-naíve cells (n=5) recorded for 25min in AFSC only.

### CRFR1 activation inhibits GABAergic transmission in females

Analysis of Stressin-1-induced changes in mIPSC frequency revealed no significant main effect of concentration of drug in females [*F*(1,42) = 0.505, *p* = 0.607], nor a significant age x concentration interaction [*F*(2,42) = 0.668, *p* = 0.518] (*Fig. 3C*). Although the main effect of age did not reach statistical significance [*F*(2,42) = 3.049, *p* = 0.081], age-specific patterns of response were observable (*Fig. 3C*) and indicated CRFR1 activation produced changes in mIPSC frequency in females; specifically, in adolescents the magnitude of decrease in mIPSC frequency became larger with increasing agonist concentrations, while adults demonstrated consistent decreases in mIPSC frequency across concentrations. Consistent with these observations, in adolescent females, only the highest Stressin-1 concentration (1 μM) significantly reduced mIPSC frequency from baseline levels [10 nM: *t*(1,7) = 1.102, *p* = 0.307, *n* = 8 cells; 100 nM: *t*(1,7) = 1.527, *p* = 0. 171, *n* = 8 cells; 1 μM: *t*(1,7) = 2.958, *p* = 0.021, *n* = 8 cells] (*Fig. 3D*). In contrast, all three concentrations of Stressin-1 produced a statistically significant reduction in mIPSC frequency in adult females [10 nM: *t*(1,7) = 3.857, *p* = 0.006, *n* = 8 cells; 100 nM: *t*(1,7) = 3.050, *p* = 0. 019, *n* = 8 cells; 1 μM: *t*(1,7) = 3.235, *p* = 0.014, *n* = 8 cells].

Analysis of mIPSC amplitude in these same cells revealed no significant main effects of age [*F*(1,43) = 0.142, *p* = 0.708] or concentration of drug [*F*(1,43) = 2.699, *p* = 0.079] in females, nor a significant age x drug interaction [*F*(2,43) = 2.366, *p* = 0.106] (*Table 2*).

### Tonic CRFR1 regulates GABAergic transmission in the CeM in an age- and sex-specific manner

To assess tonic activation of CRFR1 in the CeM, the CRFR1-selective antagonist, NBI 35965 (1 μM) was bath applied following baseline recordings of mIPSCs. There was no main effects of age within males [*t*(1,14) = 1.200, *p* = 0.250, *n* = 8 cells] or females [*t*(1,14) = 0.717, *p* = 0.485, *n* = 8 cells] in this experiment. As depicted in *Fig. 4*, within-group responses to the CRFR1 antagonist were highly variable, particularly within adolescents, and drug application did not produce overall significant changes in mIPSC frequency in either adolescent males [*t*(1,7) = 1.629, *p* = 0.147, *n* = 8 cells] or adolescent females [*t*(1,7) = 0.876, *p* = 0.410, *n* = 8 cells] (*Fig. 4*). In adults, however, CRFR1 blockade significantly reduced mIPSC frequency in males [*t*(1,7) = 5.436, *p* = 0.001, *n* = 8 cells] while producing no significant change in mIPSC frequency in females [*t*(1,7) = 0.090, *p* = 0.931, *n* = 8 cells].

**Figure 4.**
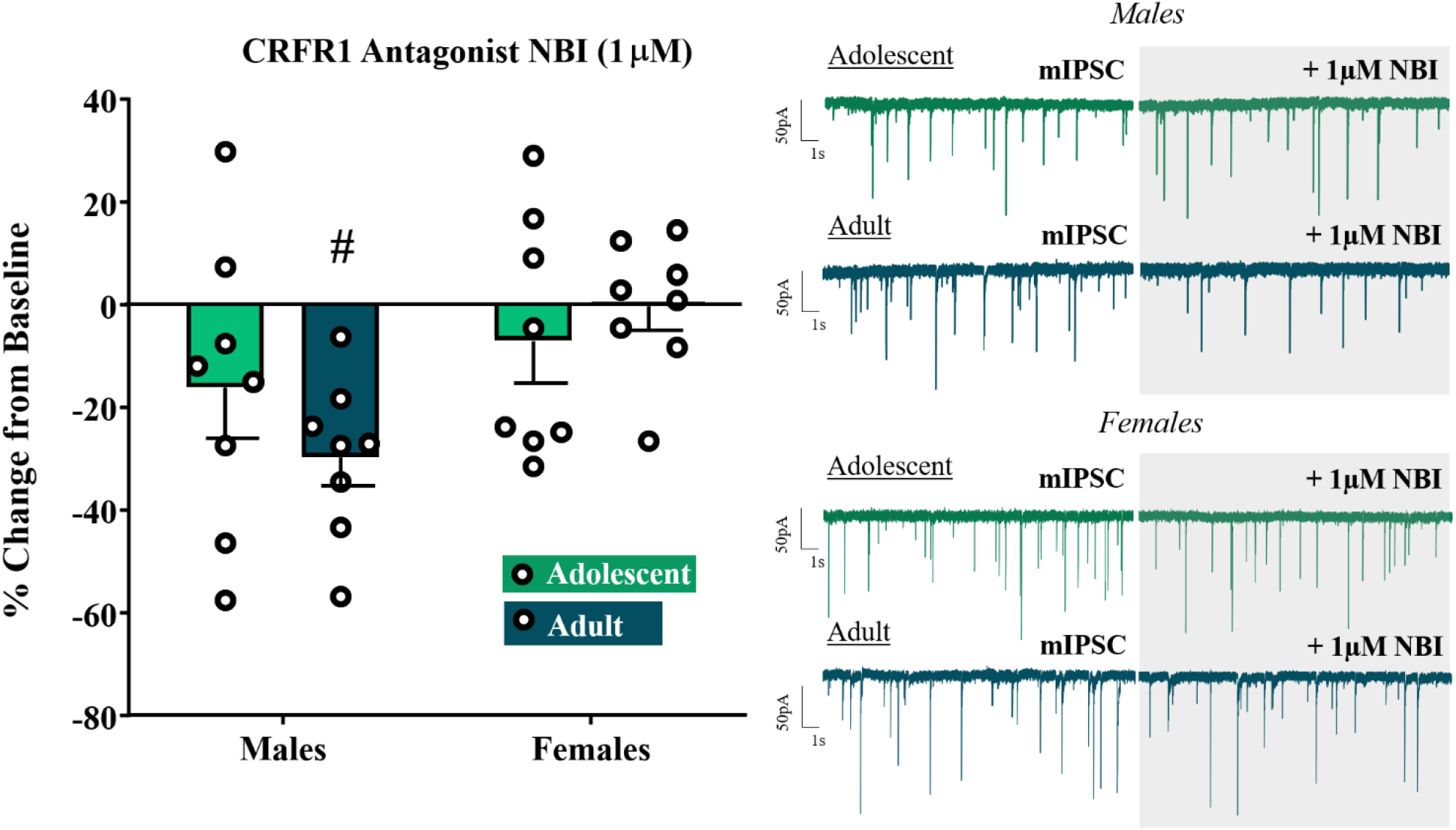
Change in mIPSC Frequency (%) following bath application of selective CRFR1-receptor antagonist, NBI (1μM). Only adult males demonstrated a significant effect of NBI. # signifies significant difference from 0.

Analysis of mIPSC amplitude in these same cells revealed no significant main effects of age in males [*t*(1,14) = 0.172, *p* = 0.866, *n* = 8 cells] or females [*t*(1,14) = 0.648, *p* = 0.528, *n* = 8 cells] in response to the CRFR1 antagonist (*Table 2*).

In a subset of adult males (*n* = 4 cells/3 animals), following completion of NBI experiments, 1 μM Stressin-1 was added to the NBI bath and recorded for an additional 10 min to verify the selectivity of the agonist. This concentration was selected because it previously produced significant potentiation of mIPSC frequency in adult males (*Fig. 3B*). In the presence of NBI, however, there were no significant changes in mIPSC frequency with the addition of the agonist [*t*(1,3) = 0.348, *p* = 0.751; Δ difference: 0.036 Hz ± 0.1035 Hz], and no significant changes in mIPSC amplitude [*t*(1,3) = 0.477, *p* = 0.666; Δ difference: 1.12 pA ± 2.348 pA].

### Length of experimental procedure impacts post-synaptic, but not presynaptic, function

Although the results of this research indicate that age and sex did not significantly influence CRFR1-selective drug effects on mIPSC amplitude, we did observe a general reduction in mIPSC amplitude across the four drug groups (three concentrations of Stressin-1 and one concentration of NBI) (*Table 2*). Importantly, these effects did not significantly differ by drug/concentration. We therefore hypothesized that observed declines in amplitude reflected a natural reduction in GABA-A receptor functioning during slice recordings under our recording conditions (averaging ~25 min length). To test this possibility, a sample of CeM neurons in adult males (*n*= 5 cells/3 animals) were patched and recorded for 30 min in ASCF only. Recordings were then assessed for run-down effects of both sIPSC frequency and amplitude.

Compared to an initial 5 min baseline recording, sIPSC frequency did not decrease after 25 min [*t*(1,5) = 1.392, *p* = 0.228; 6.230 ± 4.478% from baseline]. Importantly, however, sIPSC amplitude demonstrated a significant and notable decline after 25 min [*t*(1,5) = 3.876, *p* = 0.018; −13.88 ± 3.581% from baseline], independent of any manipulation.

## Discussion

CRF is well-established as a regulator of the physiological stress response, stimulating hormone secretion within the hypothalamus and modulating neurotransmission within extra-hypothalamic regions, including the CeM. Notably, studies incorporating diverse subject samples (i.e. females, distinct age groups) are minimal within established CRF literature, despite clinical accounts of age- and sex-specific stress responsivity in humans (Andersen, 2003; Verma et al., 2011). A recently-published review comprehensively addressed the importance of these subject factors in understanding the role of CRF system function in alcohol use disorders (Agoglia et al., 2020), and importantly, emphasized the need for future research investigating the intersections of these variables. To our knowledge, the present study is the first to directly assess how factors of age and sex interact to influence CRF’s neuromodulatory function in drug-naïve subjects. Overall, our data provide compelling evidence of a developmental shift in native and CRFR1-regulated GABAergic activity within the CeM, with distinct characterization between males and females.

Although unanticipated, our data uncovered native differences in GABAergic activity within the CeM between age groups and sexes. Adolescent females demonstrated higher sIPSC frequency than both adult females and age-matched males. Independent of age, females additionally exhibited greater post-synaptic GABA-A receptor amplitudes than males. This sex-specific receptor function could be attributed to differences in post-synaptic receptor expression, with our findings suggesting higher quantities in females than males. Alternatively, differences in GABA-A receptor subunit composition and arrangement could contribute to differential receptor functioning. Interestingly, this sex-specific GABAergic activity was not observed in action-potential independent mIPSCs; rather, in both sexes, adults exhibited greater frequencies and amplitudes than adolescents. The increase in pre- and post-synaptic GABAergic activity in adults may be indicative of developmental maturation of this system, which we have captured at an immature state in our adolescent groups. Future research incorporating younger adolescents would be beneficial for testing this theory.

In our assessment of CRFR1-regulated GABA transmission in the CeM, we uncovered both age and sex-specific regulatory function. As previously established in adult males (Herman et al., 2013b; Roberto et al., 2010), we found that CRFR1 activation significantly increased mIPSC frequency at the 1 μM concentration of Stressin-1. However, in adolescent males, CRFR1 activation *reduced* mIPSCs at this concentration, opposite of what we observed in adults. Surprisingly, this significant attenuation of mIPSCs was also observed at the 100 nM concentration, indicative of increased CRFR1-sensitivity in adolescent males compared to adults. Together, these findings may highlight an immature, hyper-sensitive function of presynaptic CRFR1 in adolescent males, and may implicate a developmental shift in this stress system. Importantly, females do not exhibit the developmental switch in CRF1R function, as both ages showed CRFR1-dependent inhibition of GABA release, with a potential increase in sensitivity in adult females. These observations in females may implicate the existence of a developmental shift in either CRFR1 function or expression, and requires future investigation.

It is possible that intracellular signaling pathways of the CRFR1 receptor are contributing to these observed age/sex effects. Although it is well-established that CRFR1 primarily couples with G-proteins and uses cyclic (c) AMP as a secondary messenger in signaling cascades, multiple intracellular signal transduction pathways have been associated with this receptor within a single brain region (Dautzenberg and Hauger, 2002). This includes Gs-coupling to stimulate adenylyl cyclase and activate PKA pathways, and G_q_-coupling to activate intracellular PKC pathways, both of which are associated with gene transcription and protein phosphorylation. Furthermore, a G-protein independent pathway has been linked with β-arrestin binding and subsequent CRFR1 downregulation (Valentino et al., 2013). Established literature hints at the importance of intracellular signaling pathways for dynamic regulation of synaptic activity; for instance, previous research in the anterior pituitary has demonstrated that sustained stress exposure can desensitize CRF-stimulated cAMP and downregulate CRFR1 (Hauger and Dautzenberg, 2000). Furthermore, evidence of CRFR1 coupling to Gi-proteins has been reported in human and rodent cells (Blank et al., 2003; Brar et al., 2004; Grammatopoulos et al., 1999; Grammatopoulos et al., 2000; Wietfeld et al., 2004), which may explain the attenuation of mIPSCs observed in our adolescent groups/adult females. Thus, shifts in proportional reliance on signaling pathways may elucidate on the age- and sex-specific responses to CRFR1 activation observed in this study.

It is worth noting that CRFR1 activation attenuated synaptic GABA release in adult females at concentrations insufficient to change mIPSC frequency in adult males. This sexually-dimorphic sensitivity has been similarly observed in the locus coeruleus, whereupon females respond to significantly lower doses of synthetic CRF than males, further supporting our conclusion that adult females demonstrate hypersensitive CRFR1 activation. Together with the opposing direction of effect observed between adult males and females, we have provided neurophysiological evidence supporting clinical reports that females and males differ in their response to stress (Kudielka and Kirschbaum, 2005; Walker and McCormick, 2009; Young et al., 2007). Of course, to determine whether these age and sex-specific physiological findings correspond to differences in the expression of anxiety-like behavior and stress response, future research is required assessing and manipulating the CeM CRF system *in vivo.*

It is important to acknowledge that the data collected from our CRFR1 experiments contained more variability within females than males. Influences of sex hormones such as estrogen could potentially account for this variability in activity (Marrocco and McEwen, 2016); however, in the few instances in which a female subject contributed two data points to the same experiment under random assignment, responses from that same female were also highly variable. This would suggest that distinct sub-populations of cells (with diverse CRFR1 function/responsivity) contribute to the variability observed in these data, more so than hormonal influences. Importantly, this variability was also similar between adolescent and adult females, reducing the likelihood of pubertal influence. However, it should be acknowledged that this adolescent age range (P40-48) represents a different stage of puberty between males and females (Bell, 2018), with an earlier onset (~P35) and conclusion (~P42) of puberty in female rats. Theoretically, as our experimental females were tested within a more mature pubertal stage than males, this could explain why distinct age differences were observed in the data collected from males but not females. To empirically test this theory, future research can investigate the CRF system within the CeM of pre-pubertal animals (~P30-33). As alluded to earlier, investigations of younger subjects would also pinpoint when shifts in CRFR1 function can be first detected, which is important for early intervention of CRFR1-targetted drug treatments for stress disorders.

Assessment of tonic CRF-regulated GABA transmission further identified age-specific differences in males, with CRFR1 blockade significantly reducing mIPSC frequency in adult males and producing no consistent change in adolescents of the same sex. In contrast, age did not influence tonic CRF-regulated GABA transmission in females. These results suggest that adult males may exhibit significantly greater CRF-CRFR1 regulated GABA release, possibly attributable to higher tonic release of CRF in this group compared to younger males and females. This is consistent with previous reports of reduced CRF mRNA in the CeA of early adolescent (P30) males and females and adult (P60) females relative to adult (P60) males (Viau et al., 2005). It is important to note that CRF peptide is both locally-generated and recruited from afferent CRF+ fibers originating from various areas, including the BNST (Schreiber and Gilpin, 2018), and the quantification of CRF concentrations within the CeM requires further targeted investigation.

When assessing cellular excitability, we determined that CeM neuronal membrane properties and resting membrane potential did not differ between age groups and sexes. However, adults demonstrated lower AP thresholds than adolescents, an effect that was most pronounced in male subjects. Furthermore, adults of both sexes demonstrated quicker response to current injection than adolescents without a difference in the amount of current injection required to produce an AP (rheobase) in either age. Finally, we discovered that when AP threshold was met, adolescent males exhibited robustly more activity than adult males, an age difference that was absent in females. These findings could be attributable to age- and sex-specific differences in voltage-gated ion channel function, although more thorough investigations would be required to make this conclusion. Alternatively, the amount of synaptic input could explain these group differences in neuronal excitability. Importantly, adolescent and adult males did not differ in sIPSC frequency, which would suggest that local phasic inhibition is not responsible for this effect. However, tonic GABA inhibition and glutamatergic input is natively abundant within the CeA and also a target of CRF (Herman et al., 2013a; Liu et al., 2004; Silberman and Winder, 2013). Since CRF acts on both inhibitory and excitatory systems, it would be worthwhile to investigate CRF1R effects on tonic GABA inhibition and the glutamatergic system within the CeM across sex and ontogeny.

Our novel and compelling findings regarding the role of the CRF system in a primary stress- and anxiety-associated brain area may underlie age and sex-specific differences in response to stressful stimuli. Established literature has positively associated CRFR1 activation with anxiogenic behavior (see review: (Schreiber and Gilpin, 2018)), in part via CRFR1-mediated increased GABA release in the CeM of adult male subjects (Herman et al., 2013b; Roberto et al., 2010), an effect we replicated. Importantly, our data suggest that CRFR1 activation *decreases* GABA release in adolescent males and females of both ages, suggesting these groups may respond to CRFR1 activation with *reduced* anxiety-like behavior. When considering age-specific stress response in humans, it is important to acknowledge that adolescents are not only phenotypically characterized with increased sensitivity to stressful situations, but also with *inappropriate* responses to anxiety-inducing events. This includes high rates of risk-taking and impaired decision-making while undergoing stress (Andersen and Teicher, 2008; Lee et al., 2003; Romeo and McEwen, 2006), behavior which has been attributed to incomplete maturation of brain regions sensitive to stress, including the amygdala (Giedd and Rapoport, 2010; Tottenham, 2017; Tottenham and Sheridan, 2009). Thus, we hypothesize that this inappropriate stress response may be attributable, in part, to age-specific CRF function within the CeM, and future investigations manipulating this system *in vivo* can elucidate on the causal relationship between CeM physiology and stress-related behaviors.

Across species, females exhibit greater susceptibility and sensitivity to stress than males, particularly within the peri-pubertal and post-pubertal window of development (see review: (Bale and Epperson, 2015)). Although our CRFR1 assessments support this characteristic hypersensitivity, the opposing direction of CRFR1-regulated GABA transmission between sexes would implicate distinct behavioral responses to stress between males and females as well. It has been previously suggested that estrogen contributes to a stress-resilient phenotype in females, an effect which corresponds to altered neurotransmitter levels within the hippocampus, frontal cortex and amygdala, although this has yet to be causally investigated. However, several additional studies have reported exactly the opposite: greater stress-induced HPA activity in females compared to males (Kudielka and Kirschbaum, 2005; Walker and McCormick, 2009; Young et al., 2007). From these studies, in combination with our own findings, we hypothesize that the stress response in females is dynamic across brain regions, and perhaps exemplary of a “stress-protective” response in extra-hypothalamic regions which seeks to compensate or correct for increased HPA axis sensitivity during development. However, future behavioral assessments will be essential for determining the sex-specific contributions of different brain regions to stress response in naïve animals.

In conclusion, this study highlights how factors of age and sex influence neurophysiological activity, and we recommend these factors be considered and reported in future investigations of the CRF system and/or CeA physiology. Furthermore, as both age and sex influence neurophysiological response to CRFR1 activation, these factors may also challenge the efficacy of CRFR1-targetted drug treatments for stress disorders in humans, and should therefore be considered for their influence in future clinical trials.

## Acknowledgments

The experiments included in this manuscript were supported by NIAAA grants P50 AA01782306, T32 AA025606 and F31 AA028166.

## Notes

Conflict of interest statement: **The authors declare no competing financial interests**.

### Competing Interest Statement

The authors have declared no competing interest.

